# Prolonged exposure to insulin causes epigenetic alteration leading to insulin resistance

**DOI:** 10.1101/2022.04.28.489884

**Authors:** Shehnaz Bano, Shyam More, Dattatray S. Mongad, Abdul Khalique, Dhiraj P. Dhotre, Manoj K. Bhat, Vasudevan Seshadri

## Abstract

Glucose homeostasis is maintained by insulin. It has been observed that hyperinsulinemia precedes insulin resistance and Type 2 diabetes. Insulin resistance is caused by multiple factors including genetic and diet. The molecular mechanism underlying insulin resistance (IR) is not completely understood. Using Glut4 and insulin receptor-expressing CHO cells we had previously shown that prolonged exposure of these cells to insulin in the absence of high levels of glucose led to insulin resistance in the cells. In the present study, we have shown that the underlying cause for the impaired GLUT4 trafficking is the defective PI3K/AKT pathway. This insulin resistance is likely due to epigenetic alterations as it is stable and can be maintained for several generations even when insulin is not provided, and epigenetic modifiers can reverse the insulin resistance. We extended these studies to liver cell line (BRL-3A) and show that these cells also develop impaired insulin signaling upon exposure to insulin in the absence of high levels of glucose. Transcriptomic analysis of the insulin-sensitive and -resistance cells uncover altered signaling networks involved in chromatin remodelling, Rho GTPases, and ubiquitination. Pathway analysis reveals the role of demethylase Kdm5b and lysine methyltransferase (Kmt2a and Kmt2e) in the development of insulin resistance. It is also observed that trimethylation of histone H3 at lysine 4 (H3K4me3) is increased in insulin resistance cellular models. We further showed that mice injected with low doses of insulin when fasting develop insulin resistance with impaired glucose tolerance and increased HOMA-IR index. Altogether, these findings suggest dysregulated synthesis of insulin in the absence of glucose stimulus could lead to epigenetic alterations that may lead to insulin resistance.

**Summary Statement:** Insulin stimulation in the absence of glucose leads to insulin resistance. We have developed a cell and mouse model of insulin resistance in this study to characterise the molecular signalling involved in insulin resistance and early onset of type 2 diabetes. The transcriptomic analysis provides new insights on epi-transcriptomic regulation in insulin resistance.

## Introduction

Insulin is the principal hormone that maintains glucose homeostasis under feeding state. This physiological process occurs through the phosphatidylinositol 3-kinase (PI3K)/AKT pathway at the cellular level^1–4^. Resistance to the metabolic actions of insulin is called as insulin resistance (IR). The conventional view of the history of diabetes suggests a reduced insulin sensitivity followed by increased insulin secretion to maintain glucose homeostasis^5,6^. Additionally, recent studies have reported an increased insulin secretion independent of IR and precede type 2 diabetes^7^. IR and hyperinsulinemia correlation makes it difficult to pinpoint actual interaction in disease state^8^. Prolonged administration of insulin causes a decreased response of insulin in diabetic patients independent of high glucose suggesting that hyperinsulinemia is self-sufficient to induce insulin resistance^9,10^.

Insulin signalling regulates glucose homeostasis mainly through activating glucose absorption in adipocytes and muscle, and by promoting glyconeogenesis in liver. In adipocytes/muscle insulin stimulates the trafficking of glucose transporter Glut4 to the plasma membrane. Previously it was shown that sensitive Chinese hamster ovary cells expressing insulin receptor and Glut4 transporter (CHO-K1) cells is insulin responsive and the insulin mediated Glut4 translocation in these cells was very similar to that seen in adipocytes^11^. It was shown that these cells traffic recombinant Glut4 to the cell membrane in insulin dependent manner and also respond to other stimuli, similar to differentiated 3T3-L1 adipocytes. Since these cells express Glut4 with Myc tag at the N-terminal region (which is exposed extracellularly) and GFP tag at the C-terminal end (intracellular), the Glut4 translocation can be analysed in these cells by FACS and microscopic methods, and have been used to understand the signalling components involved in Glut4 translocation in presence of Insulin and amino acids (^12,13,14^) or used for drug screening^15^. We have previously shown that prolonged exposure of insulin without glucose stimulus to these CHO-K1 cells expressing human insulin receptor and Glut4 transporter, leads to IR^16^. In the current study, we aimed to uncover the mechanism of IR in these cells and to assess if this IR model is also applicable to cells of hepatic origin. Insulin regulates the expression of many genes either at the transcription level or through indirect effects^17^. Global mRNA profiling has shown insulin-regulated genes in different IR models^18^. However, the interpretation is very complex due to the limitations of the existing IR model. The IR models usually describe stages preceding diabetes rather than the natural process^19^. Thus, based on the cellular model of IR we developed a mouse model of IR. We believe that such a model would closely resemble the pathophysiological alterations of cellular and molecular events of IR during the early stages of diabetes. The transcriptomic and epigenetics have not been explored much although substantial evidence has shown that they are involved in the pathogenesis of IR^20,21,22,23^. We carried out RNA-sequencing from cellular models of IR. We identified altered expression of genes involved in chromatin remodelling, mRNA splicing, transcriptional regulation, Rho GTPase and ubiquitin pathways. We investigated altered epigenome characteristics in IR state. Our studies show how transcriptomic and epigenome alterations may contribute to IR state and could define the molecular mechanism of IR. Further, we predicted a core network regulating IR by use of transcriptional and epigenomic changes associated with IR.

## Results

### Cellular IR is induced in insulin-sensitive cells in an insulin-rich environment

Insulin regulates glucose transport in muscle and adipose tissue by triggering the translocation of a facilitative glucose transporter GLUT4 from storage vesicles (GSVs) onto the plasma membrane. We used Chinese hamster ovary (CHO) cells stably expressing myc and GFP-tagged GLUT4 and insulin receptor β as a model system to investigate the effect of prolonged exposure of supra-physiological levels of insulin to skeletal muscle or adipocyte. In the basal condition myc-GLUT4-GFP is majorly sequestered intracellularly into GSVs. Upon insulin stimulation the GLUT4 translocate onto the plasma membrane and can be detected by FACS using surface myc epitope using APC channel. While the total cellular GLUT4 in the cells can be detected by GFP fluorescence using FITC channel^24^. Upon insulin stimulation the cells passaged in control medium showed an increased surface myc-GLUT4 translocation onto plasma membrane after acute insulin stimulation (Fig 1B right panel), while the cells maintained in LG (6.5 mM) + insulin (8.6 nM) medium showed a significantly reduced surface translocation of GLUT4 which was assessed by myc fluorescence of the fixed cells(Fig 1B left panel and Fig 1C). The total levels of GLUT4 in the cells were assessed by GFP fluorescence which did not change in both the cell types (Fig 1D) It suggest that the reduced surface expression of GLUT4 is not due to reduced levels of total GLUT4 among cells. This further indicates that the insulin responsiveness of cells grown in LG + Insulin condition is reduced and the cells have developed insulin resistance.

**Figure 1.**
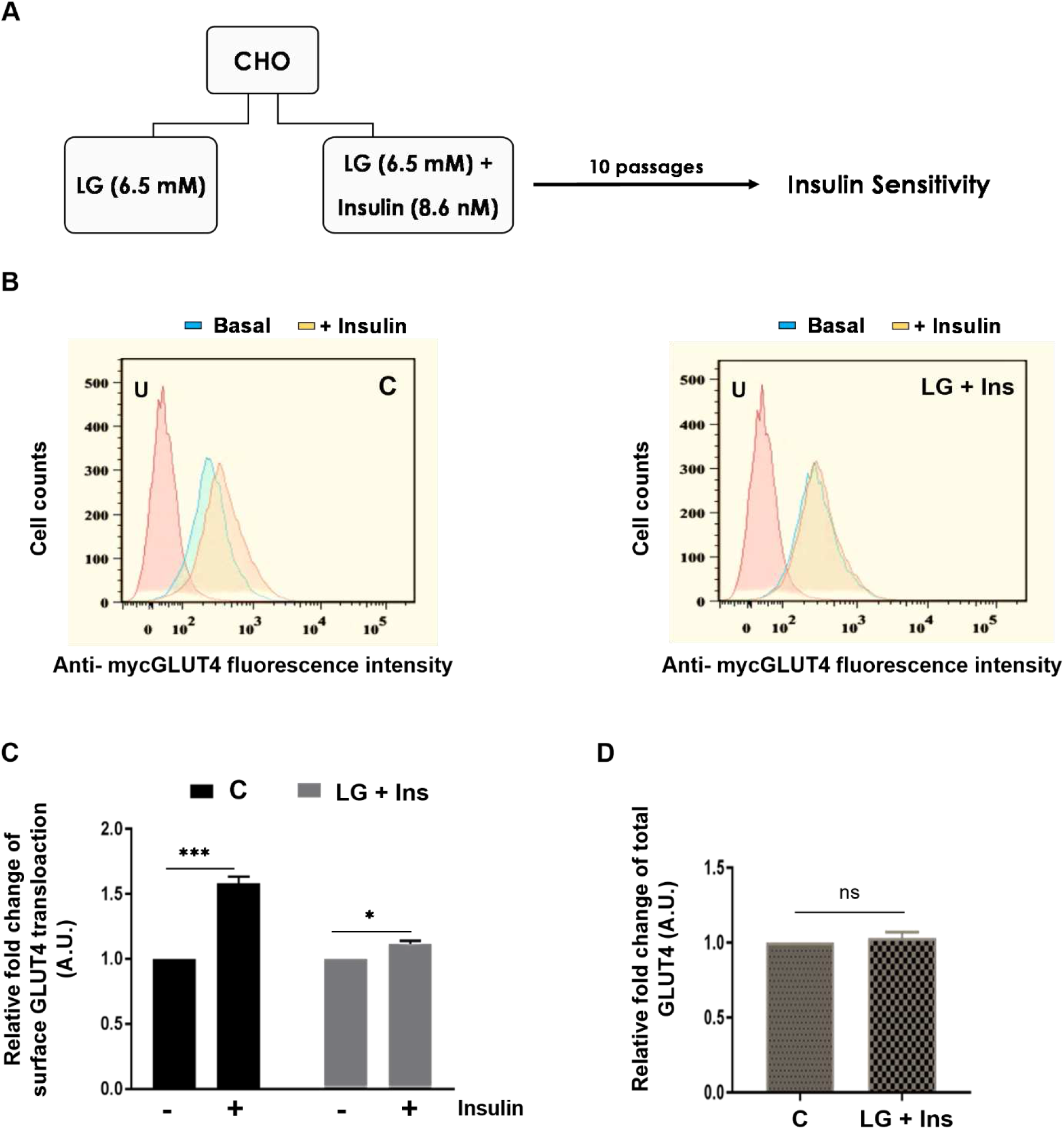
Cellular IR is induced in insulin-sensitive cells in a diabetic environment. **A:** Schematic representation of workflow used to establish insulin resistance. **B:** GLUT4-CHO cells were fixed and stained with anti-myc mouse monoclonal antibody without permeabilization to monitor the membrane expression of GLUT4. The Fluorescence intensity for the population of cell maintained in **(B Left panel)** low glucose medium and **(B Right panel)** low glucose media with insulin was measured. Mean fluorescence intensity (MFI) was calculated using FlowJo software (V10). **C:** Relative MFI was calculated by normalizing with no insulin MFI and graph was plotted between relative fold change upon insulin stimulation for cell population that were grown in indicated medium. **D:** GLUT4-CHO cells were permeabilized and FITC channel was used to monitor the total expression in cell against GFP-GLUT4. The Fluorescence intensity for the C cells and IR cells were measured. Mean fluorescence intensity (MFI) was calculated using FlowJo software (V10). Relative fold change was calculated by normalizing with C cells and graph was plotted. The data represents mean±SEM. *P < 0.05, **P < 0.01, ***P < 0.001, ****P < 0.0001 (n=3); ns means non-significant P>0.05. U represents unstained cells.

### Insulin resistance state is defined epigenetically

To determine the nature of insulin resistance i.e., temporary or a permanent change, IR cells were grown in low or high glucose medium like C cells without the presence of insulin for several passages (5-10 passages) and then the cells were assessed for insulin signalling. We observed that the IR cells were showing resistance even after maintaining them in normal growth medium for several generations. These cells did not show increased insulin stimulated GLUT4 translocation and remained insulin resistant (Fig 2A). This result suggests that the defective insulin signalling in these cells might be due to an epigenetic change that are stably maintained through somatic division.

**Figure 2.**
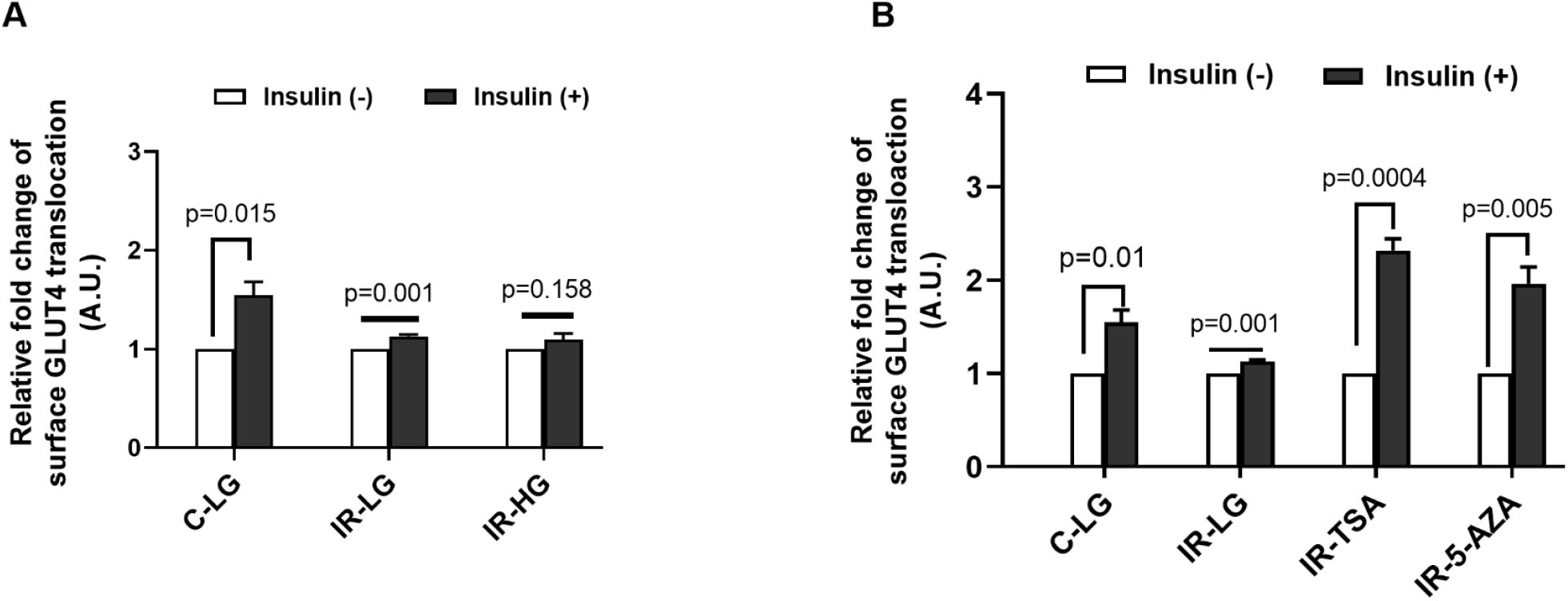
IR state is defined epigenetically. **A:** Insulin stimulated GLUT4 translocation to plasma membrane of C and IR cells when grown in low glucose (LG) and high glucose (HG) medium for ten to five generations respectively. Relative fold change was calculated by normalizing without insulin MFI and graph was plotted between relative fold change upon insulin stimulation using flow cytometry. **B:** Insulin responsiveness of GLUT4 translocation to plasma membrane of IR cells when grown in TSA and 5-AZA treated in low glucose medium was examined. Relative fold change was calculated by normalizing without insulin. MFI and the graph plotted for relative fold changes upon insulin stimulation using flow cytometry. The data represents mean±SEM; n=3.

If IR is due to epigenetic changes, then the treatment of cells with epigenetic modifiers may be able to reverse the insulin resistance phenotype. 5-aza-deoxycytidine (5-AZA) and trichostatin A (TSA) are two epigenetic modifiers that have been used for restoring insulin sensitivity of IR cells^25^. Therefore, we treated the IR cells with these epigenetic modifiers to reverse the resistance phenotype. We have grown IR cells in the presence of these epigenetic modifiers in growth medium for 24 hours and assessed insulin sensitivity of these cells. We found that the insulin stimulated GLUT4 translocation was significantly increased in the IR cells treated with epigenetic modifiers (Fig. 2B). This suggests that the IR is maintained due to epigenetic alterations and the resistance phenotype can be reversed by epigenetic modifiers.

### Impaired PI3K/AKT signaling in insulin resistant CHO-GLUT4 cells

Insulin stimulation leads to activation of a series of downstream signals through binding to its receptors^26,27^. In this process, PI3K activation consequently activates AKT by phosphorylating at position Serine (S) 473 and threonine (T) 308. This AKT phosphorylations lead to increased membrane localisation of GLUT4, which facilitates glucose uptake in adipose tissue and skeletal muscle. Any defects or abnormalities in insulin related signaling pathways may lead to IR ^28,29^. We evaluated the insulin stimulated PI3K/AKT pathway activation in the insulin - sensitive and -resistant CHO cells. We examined insulin stimulated phosphorylation of AKT at position Serine (S) 473 and threonine (T) 308 in these cells. There was a reduced insulin-stimulated phosphorylation of AKT at position S-473 & T-308 without significant change in the quantity of total Akt protein in IR cells compared to C (Fig. 3A, B, C and D). These results suggests that the IR may be due to deregulated PI3K/Akt signaling in response to insulin.

**Figure 3.**
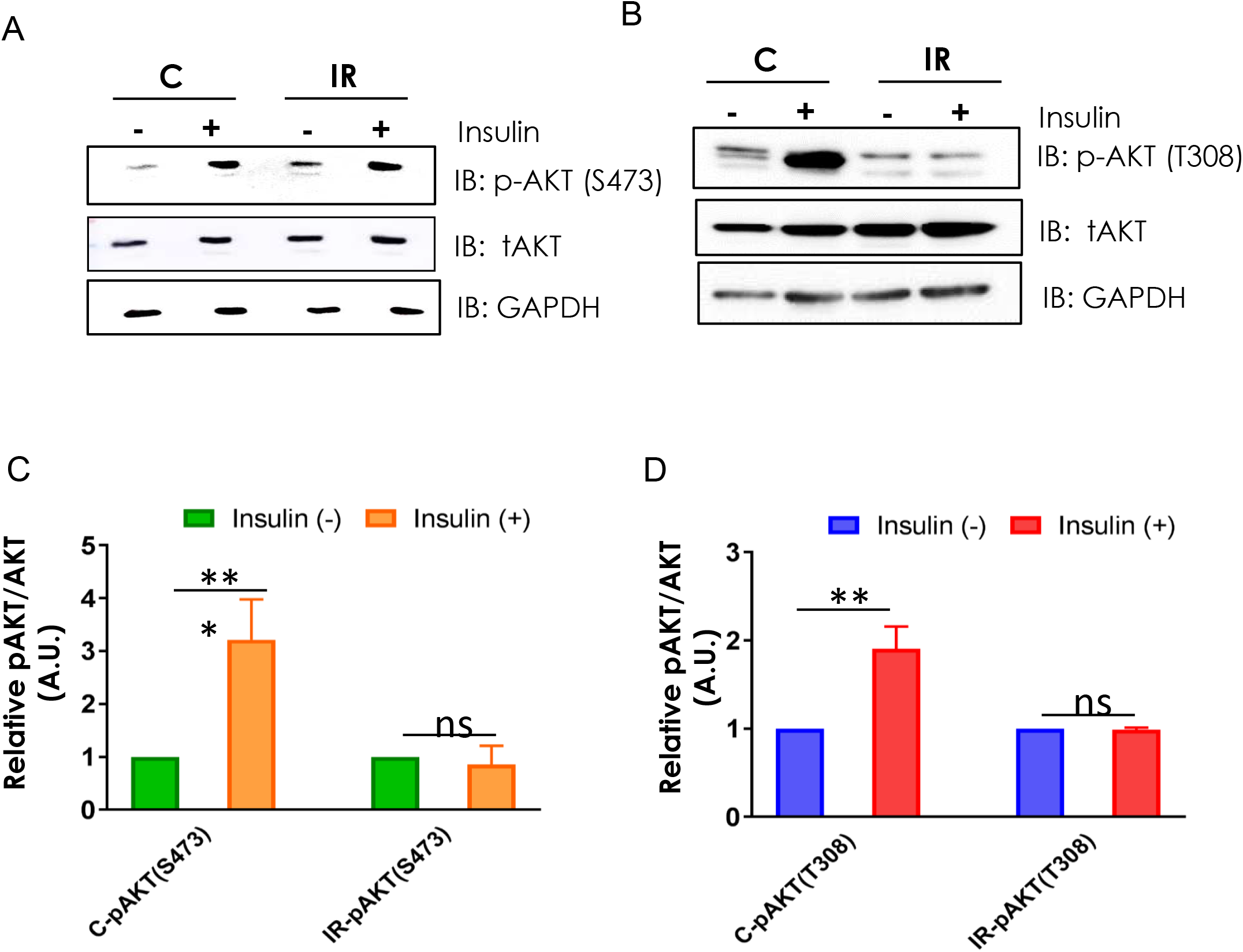
Prolonged insulin exposure ameliorates insulin signaling through PI3K-AKT signaling. **A-B**: The phosphorylation of Akt was assessed in IR and C cells by probing for phosphorylated Akt, total Akt and GAPDH**. C-D**: Densitometry for the is shown, and data are expressed as arbitrary units in relation to the basal of the same. Protein quantification was adjusted for the corresponding GAPDH level. The data represents mean ± SEM. *P < 0.05, **P < 0.01, ***P < 0.001, ****P < 0.0001 (n=3)

### Hepatic origin cells exposed to insulin in the absence of excess glucose become IR

Insulin signalling is conserved across cell types even though the ultimate output of the signalling varies in peripheral tissues. In skeletal muscle and adipocytes insulin signalling results in GLUT4 translocation leading to increased glucose uptake. While in hepatocytes, it results in activation of glycogen synthesis and glucose uptake is not significantly affected. Together, the final output is different in insulin target tissues, but PI3K/AKT signalling is the principal insulin signaling in all cell types. Since we have shown that the IR is due to the defective PI3K/AKT signalling, therefore, we ask the question whether hepatocytes BRL-3A cells would become IR using similar strategy. We have assessed the effect of insulin exposure in the absence of high levels of glucose on BRL-3A hepatocytes. We maintained BRL-3A cells for about 10 passages in control medium or LG (6.5 mM) + Ins (8.6 nM) medium. We determined the regulation of GSK3 activity in LG + Ins cells and control (C) cells upon insulin stimulation. We observed that insulin stimulation does not significantly increase the phosphorylation of GSK3β and AKT (S473) in case of LG + Ins cells while there was a robust stimulation of GSK3β and AKT-S473 phosphorylation in C cells (Fig. 4 A, B, C and D). We could find attenuated insulin signalling in hepatocytes. This supports the fact that the prolonged exposure of cells to insulin in low glucose growth medium causes IR in peripheral tissues.

**Figure 4.**
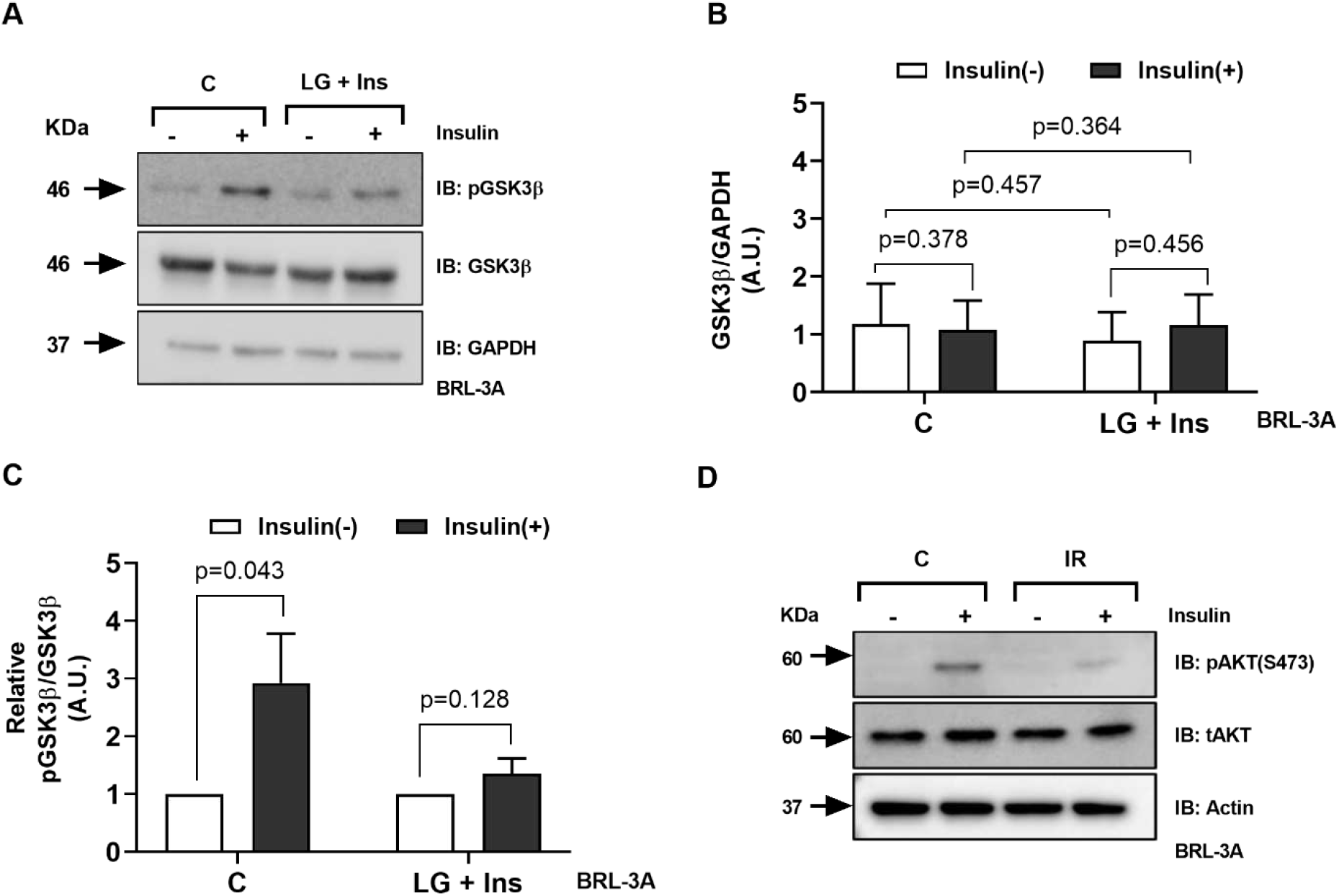
Hepatic origin cells (BRL-3A) exposed to insulin in the absence of excess glucose become insulin resistant. **A:** The levels of phosphorylated GSK3β, total GSK3β and loading control GAPDH was assessed by wester blotting in extracts prepared from cells passaged for ten generations in low glucose (C) and low glucose with insulin growth medium (LG + Insulin) (IR). Densitometry analysis of the band intensity was performed and the ratio of GSK3β to GAPDH (**B**) and ratio of Phosphorylated GSK3β to total GSK3β (**C**) is shown. The ratio is of band intensity expressed in arbitrary units. Protein levels were normalised to corresponding GAPDH level. The data represents mean±SEM. *P < 0.05, **P < 0.01, ***P < 0.001, ****P < 0.0001 (n=3); ns means non-significant. **D:** The phosphorylation of Akt was assessed in extracts prepared from cells passaged for ten generations in low glucose (C) and low glucose with insulin growth medium (LG + Insulin) by probing for phosphorylated Akt, total Akt and GAPDH.

### Diverse transcriptional changes associated with cellular IR models

To understand the molecular mechanism by which IR was induced, we performed transcriptomic profiling of the IR cellular model with respective C. The genes deregulated in hepatocytes BRL-3A cells are shown in heatmap form, volcano plot, KEGG pathway and reactome pathway (Fig. 5 A, B, C and E). To find out the related functions of the differentially expressed transcripts in the IR cells, we performed Gene ontology (GO) analysis which classify gene function based on molecular, cellular, and biological processes (Fig. S1). To further examine how deregulated transcripts coordinated with respective target genes to regulate insulin signaling. We constructed protein-protein interaction networks from gene expression data using string analysis, to better visualise the effects of IR at a system level. This approach identified a set of genes that were highly coordinated in their response to insulin signaling, with some included in a single pathway (Fig 5D). Implying a potential role of differential expressed RNAs in IR. The IR induces impairment either directly or indirectly affecting insulin signaling, thereby affects glucose homeostasis^30^. To further examine the molecular mechanism of IR we correlated dysregulated gene function effects on insulin signaling. We also performed one pilot study of RNA-Seq in CHO-GLUT4 cells to identify conserved pathway dysregulated in our cellular IR models, we intersected gene expression datasets from BRL-3A and CHO cells. We identified PI3K/AKT signaling, chromatin remodelling, transcriptional regulation, ubiquitination and proteasome degradation and Rho signal transduction are commonly affected in both systems (Fig 6 and S2). Together data from BRL-3A and CHO cells showed differential expression of genes involved in histone methylation activation. Such as in CHO cells we found upregulated expression of Kmt2a gene. While in BRL-3A cells we found upregulated expression of Kmt2e and Kdm4b gene along with downregulation of Kdm5b gene. In general, Kmt2a and Kmt2e are methyl transferase and promote histone H3K4 methylation, majorly H3K4me3. While Kdm4b and Kdm5b are histone demethylase. Interestingly Kdm5b inhibits methylation of H3K4 while Kdm4b promotes methylation of H3K4me. Together, it supports the notion that H3K4me3 might play a role in insulin resistance.

**Figure 5.**
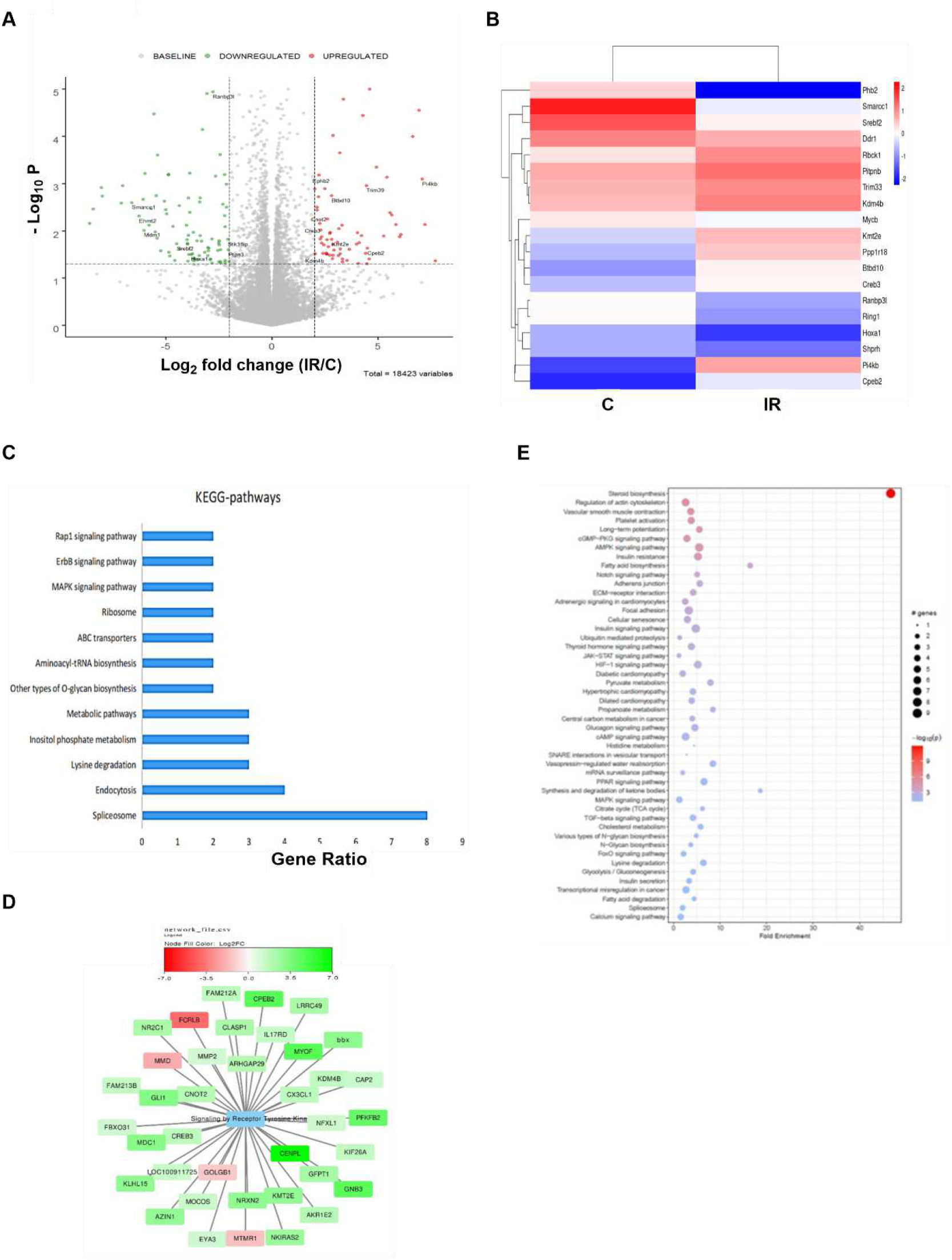
Diverse transcriptional changes associated with IR hepatocytes (BRL-3A) cells. **A:** Volcano plot illustrating fold change (-log10 adjusted P-values) in expression for all genes in cells. **B:** Heat map of the differentially expressed genes. **C:** KEGG pathway enrichment in IR vs C cells. **D:** Chart representing differentially expressed genes enriched under reactome pathway: signaling by tyrosine kinase. **E:** Chart indicating the top reactome pathways of differential genes, among the IR cells vs. C cells of BRL-3A.

**Figure 6.**
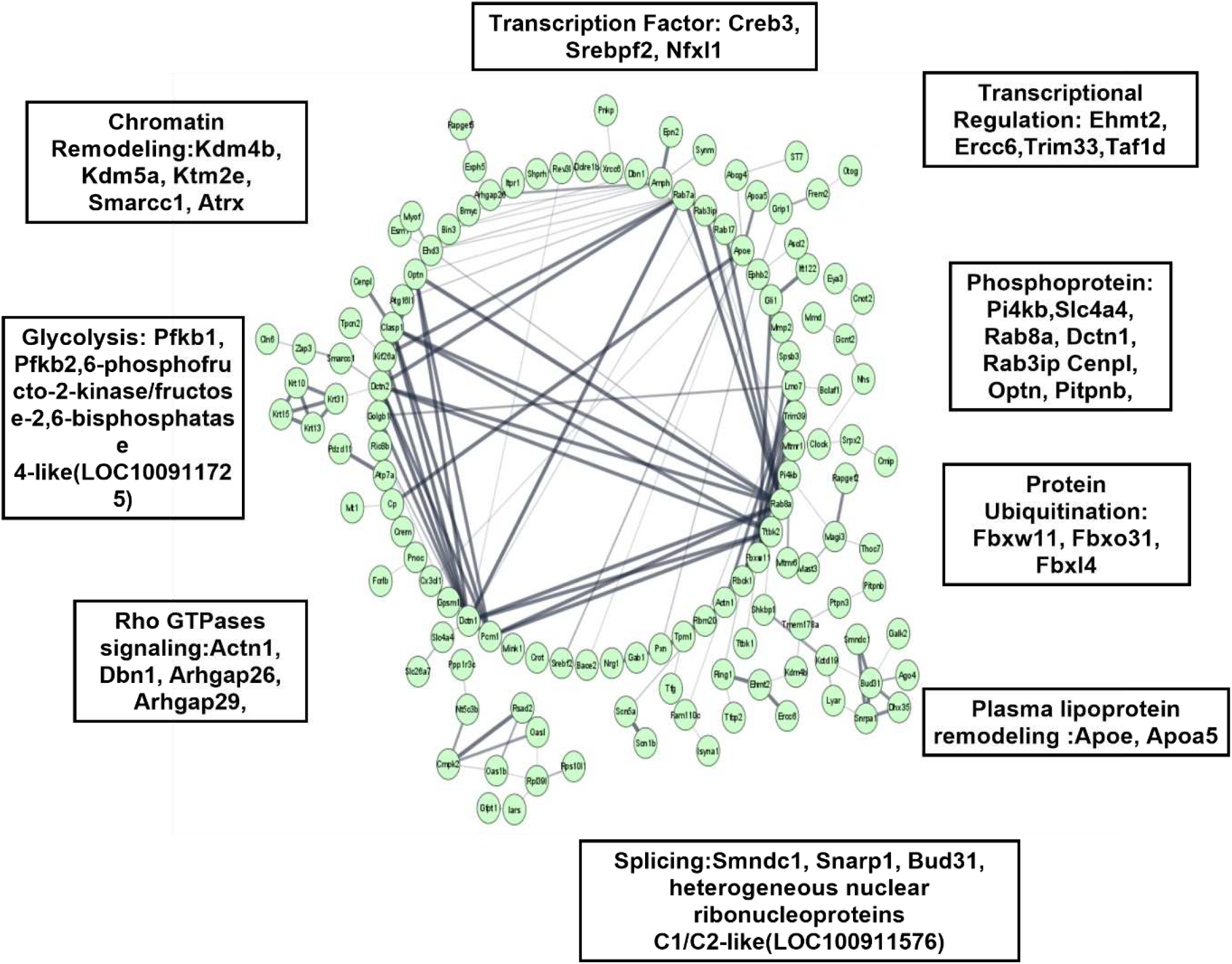
Protein protein interactions associated with IR. Network of predicted protein-protein interactions from STRING analysis (Szklarczyk et al., 2017) using insulin resistance regulated genes in BRL-3A IR cells.

### Elevated H3K4me3 is associated with IR

Our results suggest that epigenetic changes are associated with IR, which if reversed may restore insulin sensitivity. We investigated whether there was a relationship between IR and genome wide methylation of histone H3 in IR cellular models. We analysed the extent of Histone 3 specific methylation using site- and modification-specific antibody in insulin resistant and insulin sensitive CHO and BRL-3A cells. The levels of H3K4me3 are increased in IR cells compared with control cell (Fig. 7 A, B, C and D).

**Figure 7.**
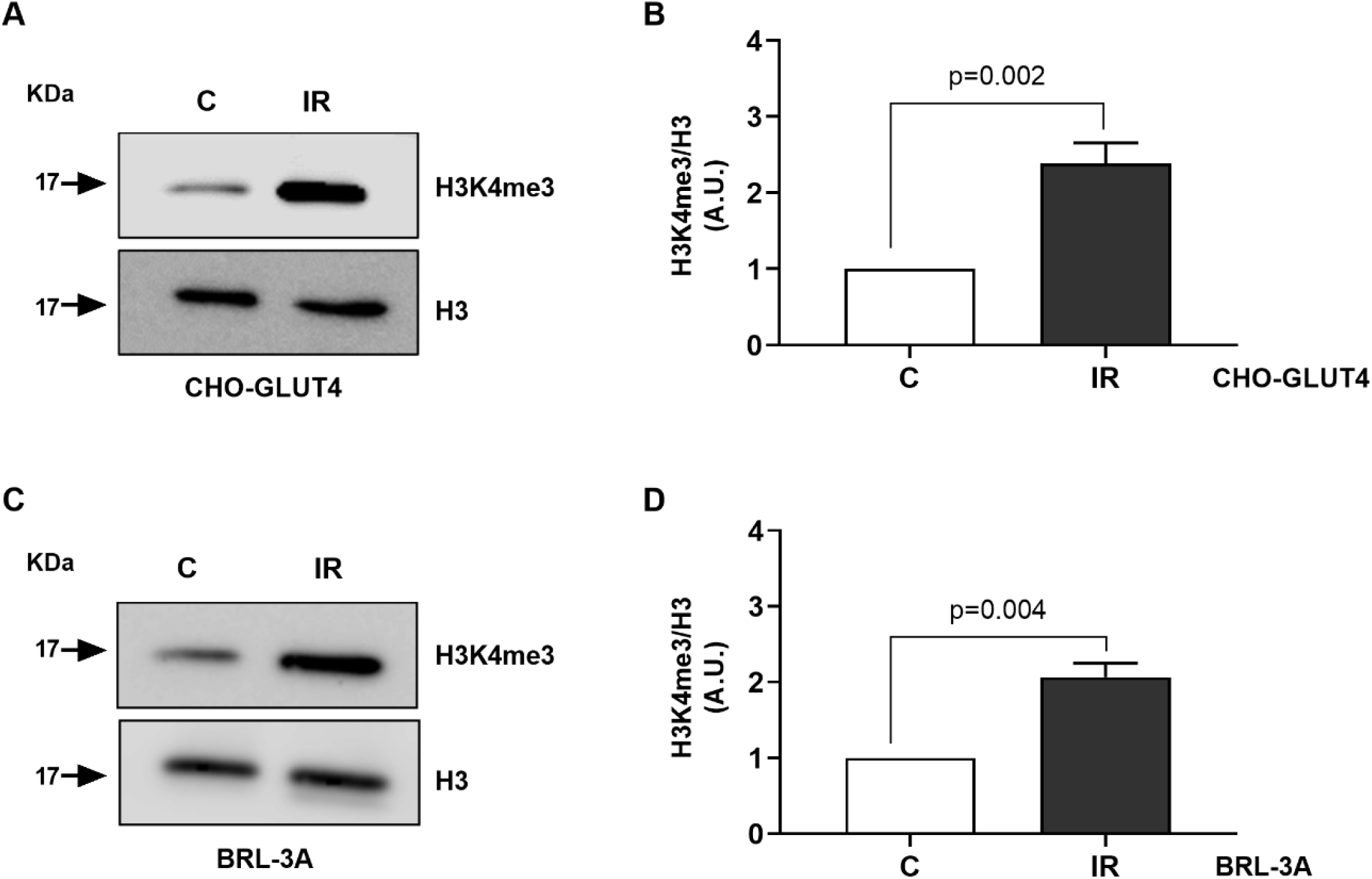
Elevated H3K4me3 is associated with IR. Western blotting for trimethylated H3 (K4) and total H3 in insulin sensitive and resistant CHO (**A**) and BRL-3A (**C**) cells. The band intensity was assessed by densitometric scans and the ratio of trimethylated H3 (K4) to total H3 was plotted with control cell ratio set at 1 (**B** and **D** respectively). The band intensity was measured in arbitrary units and the data are expressed as mean of three biological repeats.

### Mouse model of IR

Since our results suggested that exposure of cells to insulin in the absence of high level of glucose lead to IR in insulin responsive cells, we wanted to explore if the same can be extended to an animal by exposing the mice to insulin in the absence of high levels of glucose. Briefly, we injected defined amount of insulin (90 mU/Kg) every day to fasting mice and let the mice continue fasting for further 2 hours for about 15 days. We believe that exposure of animal to insulin in the absence of glucose (fasting) simulates the conditions of LG + insulin for the cells. We do not see any effect of fasting and insulin injection alone on glucose tolerance as shown in Fig S3C. Therefore, for the main experiment we have used two groups: 1) C and 2) LG + Ins (Fig.8 A). After 15 days we measured fasting glucose and performed GTT assay. We observed that the clearance of injected glucose in time frame of 2-h was delayed in LG + Ins group compared to C group (Fig. 8 B), indicating that LG (fasting) + Ins group had impaired glucose homeostasis. Consistent with this impairment, area under curve (AUC) of blood glucose response were significantly elevated in LG (fasting) + Ins group relative to control group (Fig. 8 C). We further tested the insulin sensitivity in both the groups using an ipITT. We found that insulin induced hypoglycaemic response was impaired in LG + Ins group, as their blood glucose levels were elevated following intraperitoneal insulin (Fig. 8 D). Fasting plasma insulin, and glucose levels were significantly higher in LG (fasting) + Ins group compared with control group (Fig. 8 E and 8F). Despite the elevated levels of insulin, blood glucose was still not cleared in LG + Ins group, indicating that these mice have developed IR. We found that HOMA-IR index is significantly increased in LG + In compared to C group (Fig. 8 G). We have also examined percentage body weight in both the groups to determine obesity at the end of the experiment in IR model (Fig. S3C) and found no significant differences amongst the groups at the end of the experiment. Overall, it suggests that LG (fasting) + Ins group has developed IR and shows impaired insulin signaling. Furthermore, the results of H&E staining revealed that no obvious tissue damage, such as inflammation and/or necrosis, was observed in tissue sections of the liver, white adipose tissue, and skeletal muscle (Fig. S3D), which indicated that the IR is at very early stages. We observed significantly enlarged volume of pancreatic islets and adipose tissue (Fig. 8 H and I). Together, this IR model demonstrates the early phase of type 2 diabetes and might be a good model to study IR.

**Figure 8.**
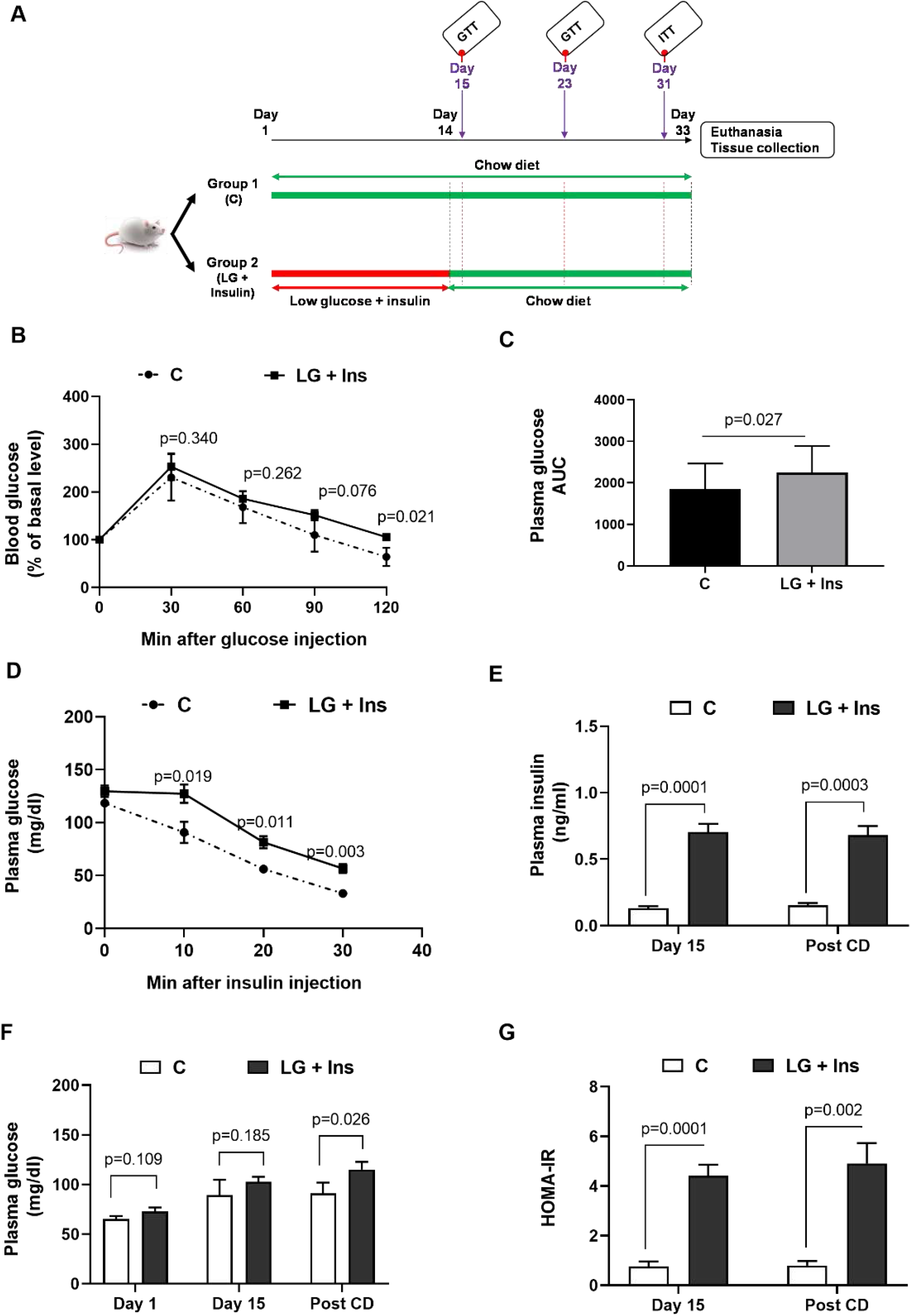
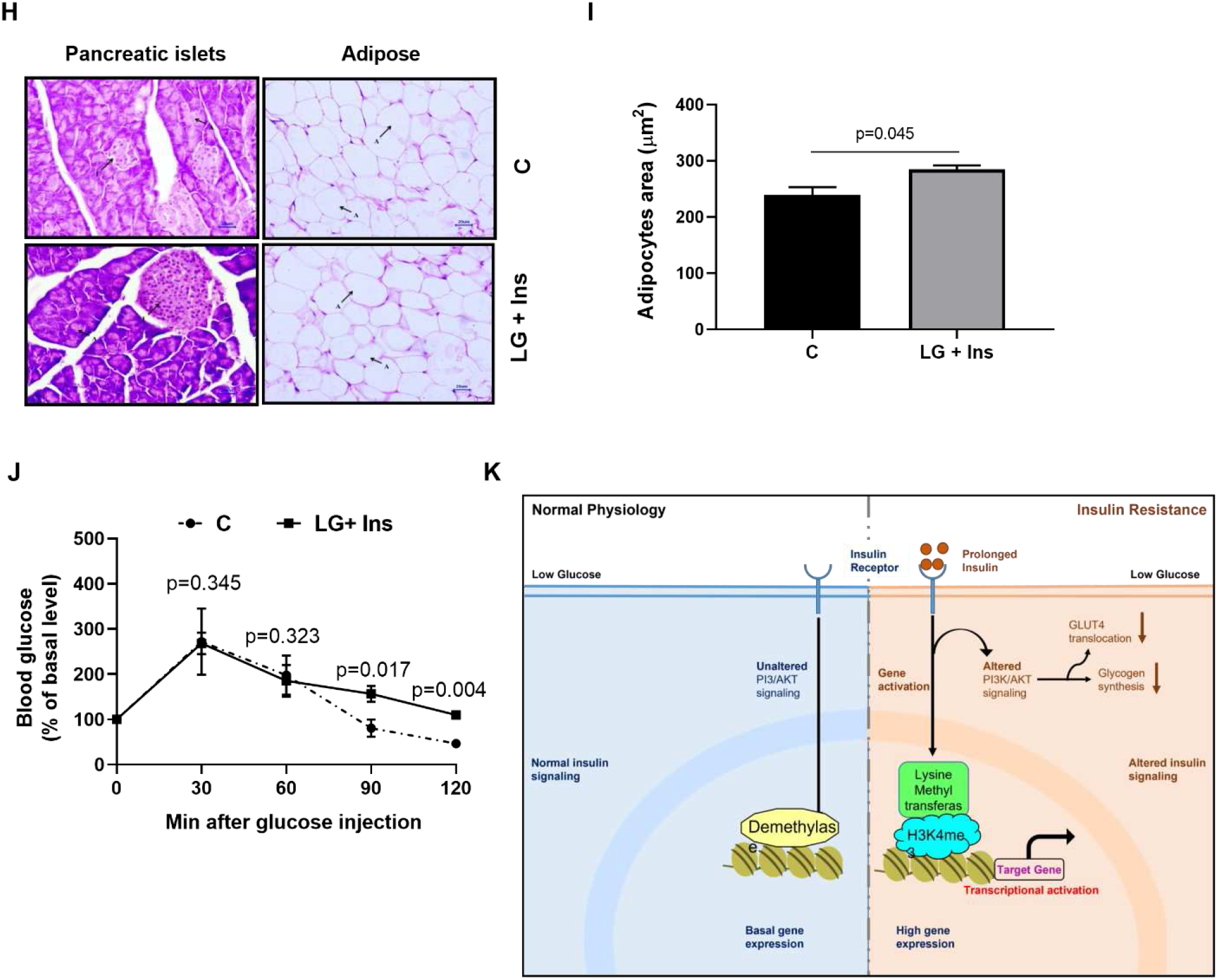
Mouse model of IR. **A:** Schematic workflow of development of mouse model of IR. **B:** Insulin sensitivity assay by glucose tolerance test after two weeks of post treatment: Plasma glucose concentrations during the intraperitoneal glucose test (IPGTT;3g/kg body weight) following 8 h of fasting in Swiss Albino male mice were estimated at various time points and graph was plotted for the same. The data represents mean±SEM; C: n=3; LG + Ins: n=6. **C:** Area under the curve (AUC) of blood glucose levels. **D:** Insulin sensitivity assay by insulin tolerance test after post removal of treatment. For ITT, plasma glucose concentrations were measured at different time points using hand-held glucometer during the intraperitoneal insulin injection (0.25U/Kg body weight) after 2 h of fasting in Swiss Albino male mice. The graph was plotted for ITT. The data represents mean±SEM; C: n=3; LG + Ins: n=6. **E-F:** The fasting insulin and fasting plasma glucose was measured at different days using glucometer and insulin ELISA kit respectively and the graph was plotted for the same. The data represents mean±SEM; C: n=4; LG + Ins: n=6. **G:** Insulin sensitivity assay by HOMA-IR index was calculated using fasting glucose and fasting insulin values and graph was plotted. The data represents mean±SEM; C: n=4; LG + Ins: n=6. **H:** Haematoxylin and eosin staining from mice tissue. All images are presented at a magnification of x400. The upper and lower panel represent histology of tissue taken from C and LG + Ins group, respectively; where islets and pancreatic acini are indicated in the pancreas section; while adipose tissue is indicated in the adipose section. **I:** Adipocyte area was calculated using image J, from 100 adipocytes of each group. The data represents mean±SEM; n=3. **J:** Blood glucose levels in response to glucose load was assessed by glucose tolerance test after one week of no treatment: Plasma glucose concentrations were estimated at various time points and graph was plotted for the same. The data represents mean±SEM; C: n=3; LG + Ins: n=6. **K:** Graphic summary of our current model. Under basal condition, there is low expression of genes to maintain basal glucose homeostasis. During prolonged insulin exposure in low glucose condition, activation of insulin receptor sustains basal AKT activation. Due to which downstream signaling is not further activated. This prolonged insulin exposure leads to recruitment of lysine methyl transferase to the histone H3 followed by tri-methylation at K4 thereby inducing activation of target gene promoter. This activation eventually altered insulin signaling by modulating transcripts levels and ultimately leads to insulin resistance.

To further characterise whether this induced IR is stable or not due to altered diet/fasting we examined the glucose homeostasis in IR group using GTT, again after a week of keeping them on continuous accesses of normal chow diet feed like control group and with no insulin injection. We observed that the blood glucose levels were significantly elevated in LG + Ins group compared to control groups (Fig. 8 J). We found that the resistance persisted in IR group, suggesting that the altered insulin sensitivity is stable and may be due to changes at epigenetic level.

## Discussion

The aim of present study was to understand the mechanism of how prolonged exposure of physiological levels of insulin in the absence of glucose stimulus induced insulin resistance. Tabrak et al., 2009 has shown that insulin resistance is a slow progressive disorder^31^. It results in abrupt increase of fasting glucose even in the absence of glucose stimulus, just before the diagnosis of type 2 diabetes. Therefore, in our early findings we established insulin resistance model to test this speculation. Here we found that our *in vitro* IR models mirror many of the defects seen in hyperinsulinemia, thus providing a unique disease-in-a dish model to study the molecular alterations underlying these abnormalities. Exploiting these models, we have observed the underlying cause is conserved across cell type. The IR was not reversed by growing cells without insulin for several generations, reflecting the root might be due to altered epigenetics in IR state. Indeed, epigenetic modifiers 5-aza and TSA could rescue insulin sensitivity of IR cells. Both drugs could influence stability of methylation and/or chromatin marks either directly or through their modifying proteins, implying a converging mechanism to IR. The underlying cause of IR defines a novel signature of cell autonomous alterations due to lack of multiple interference like high glucose, high insulin, lipid, and hormones. Furthermore, loss of transcriptional integrity is linked to uncontrolled diabetes. Our data from RNA-sequencing revealed that the prolonged insulin exposure regulates a multi-dimensional network of gene expression in insulin signaling. Earlier studies suggested that the fasting regimes can be more effective for improving insulin sensitivity. The prolonged exposure of physiological levels of insulin under hypoglycaemia revealed non-beneficial effects of low glucose. Our results could identify novel genes that are reprogramed by prolonged insulin exposure in the absence of effective insulin signalling. We identified many genes which directly or indirectly regulate histone methylation, specifically H3K4 trimethylation showed increased expression to both cellular IR models. It confirmed that H3K4me3 has a role to play either at transcription or mediating insulin resistance signaling with other regulatory inputs in environmental-context dependency. Later, we found that H3K4me3 is positively associated with insulin resistance in humans^32^.Further work is also required to understand the interplay between H3K4me3 and insulin signaling to maintain glucose homeostasis. Based on the above results, the model is proposed on how molecular mechanism of insulin resistance was dysregulated compared to insulin sensitive state in Fig 8 K. In the present study, we established a new mouse model to study IR. The protocol was based on mimicking physiological state of pre-diabetes subjects in urban lifestyle. Together, the results confirmed that the IR mouse model resembles the phenotypes seen in pre-diabetes patients. We believe that this model can be used to study defects observed at an early stage of type 2 diabetes.

## Research Design and Methods

### Cell culture

Chinese hamster ovary (CHO) cell line expressing human insulin receptor and glucose transporter 4 (with myc and green fluorescent protein at N- and C-terminal respectively) was kindly provided by Dr. Manoj K. Bhat (NCCS, India). Rat liver cell line BRL-3A was obtained from the cell repository of the National Centre for Cell Science (NCCS). The Glut4-CHO and BRL-3A cells were maintained in 16.5 mM mannitol containing DMEM-6.5 mM low glucose (LG) medium supplemented with 10% fetal bovine serum, 100 units/ml penicillin and 100 μg/ml streptomycin. Cells were maintained in a humidified 5% CO_2_ incubator at 37°C.

### Cellular model of IR

Cells were grown in DMEM-LG medium with 8.6 nM of insulin (LG + Ins) and later insulin sensitivity of these cells was assessed. Cells were treated with 8.6 nM insulin for 10 min to give acute insulin stimulation.

### Flow cytometry

To quantitate the surface GLUT4 onto plasma membrane, we followed FACS protocol mentioned in our earlier article (11). The total GLUT4 was quantified by recording GFP fluorescence. The parameters were recorded for ten thousand events for each experimental condition. Samples were acquired on FACSCanto™ II analyzer (Beckton Dickinson Biosciences) using the FACSDiva™ software. The acquired data (.fcs files) were further analyzed using FlowJo® software (FlowJo, LLC).

### Epigenetic modifiers treatment

IR CHO cells (2 x 10^6^) were seeded in a complete DMEM-LG medium. After 24 h, the medium was removed, and cells were pre-treated with either 5 μM 5-aza-deoxycytidine (5-AZA) and 200 nM trichostatin A (TSA) in fresh DMEM-LG complete medium for 24 hours. The respective solvent was used as a control treatment. Following 5-AZA and TSA treatment, the cells were further examined for insulin sensitivity.

### Immunoblotting

Cells (3*10^6^) were washed with ice-cold PBS and lysed in RIPA buffer on ice. Lysates were cleared by centrifugation at 12,000 rpm for 15 min at 4°C. The supernatant was used as a whole-cell protein lysate. Protein concentration was quantified by Bradford assay with Coomassie Plus Protein Assay Reagent (Thermo Fisher Scientific, CA, USA).

### Transcriptomic sequencing and analysis

The TRIzol homogenates were sent to the service provider (Bionivid/Nucleome). The quality of RNA was assessed using a bioanalyzer (Agilent); only RNA with a RIN >8 was used for library preparation. Libraries were sequenced on an Illumina Nova-Seq/Hiseq instrument in a pair-end, 150-bp (PE150) mode. The paired-end reads were filtered using fastp tool with Phred score cut-off of >Q30.^33^. Rattus norvegicus Reference Genome Rnor_6.0 downloaded from Ensembl (GCA_000001895.4) was used as reference for mapping of reads and identification of transcripts for BRL-3A cells. While cricetulus griseus reference genome from Ensembl was used for CHO-GLUT4 cells reads. Hisat2 pipeline^34^ was used for reads alignment and transcripts were identified using Stringtie^35^, using default parameters for both the tools. Differentially expressed transcripts between control and treated samples were identified by DESeq2^36^ using a fold-change threshold of absolute fold-change >=1.0 and a statistically significant Student’s t-test *P* value threshold adjusted for false discovery rate of less than 0.001. Unsupervised hierarchical clustering of differentially expressed genes was done using Cluster R package (v3.0) and visualized using Java Tree View (http://jtreeview.sourceforge.net/). Gene ontologies and KEGG pathways that harbour expressed transcripts were enriched using DAVID Functional Annotation Tool [DAVID Bioinformatics Resources 6.8, NIAID/NIH^36^. ^36^Statistically significantly enriched functional classes with a *P* value adjusted for false discovery rate of less than 0.05 derived using the hypergeometric distribution test corresponding to differentially expressed genes were determined using Student’s t-test with Benjamini Hocheberg FDR test^37^. All the visual representation were made using R packages. For Heatmaps ggplots was used, EnhancedVolcano was used to generate the volcano plots. Venny (https://bioinfogp.cnb.csic.es/tools/venny/) as used to generate the venn diagrams. For Networks analysis a comprehensive network visual analytics platform for gene expression analysis named Network analyst was used. The Reactome pathways were enriched using clusterProfiler^38^ R package and network was plotted using Cytoscape^39^.

### Mouse IR model

Swiss albino male mice of 8-10 weeks’ old were allocated from the Experimental Animal Facility (EAF) at NCCS and acclimatized for one week. The mice were housed and maintained in animal quarters under environmentally controlled conditions (22 ± 2°C) with a 12 h light/dark cycle. They had free access to water and standard rodent pellet food (Normal chow diet) *ad libitum*. All animal experiments were performed as per the requirement and guidelines of the Committee for the Purpose of Control and Supervision of Experiments on Animals (CPCSEA), Government of India, and after obtaining permission of the Institutional Animal Ethics Committee (IAEC). The mice were randomly segregated into four groups.

Here we have designed fasting condition in such a way to avoid the effect of severe fasting on insulin signaling. We have also tried day fasting on mice, but we do not see any effect on insulin signaling due to nocturnal nature of mice (Data not shown here). Therefore,to develop insulin resistance, mice were kept on ten hours of fasting to mimic low glucose condition (LG) at 12:am-10am. And an intraperitoneal insulin (Ins) injection of 8.6 nM in 0.9% saline solution was given and kept for further fasting for two hours. The mice were kept further for two hours of fasting after insulin injection to understand the effect of supra physiological levels of insulin onto insulin response of peripheral tissues in low glucose condition in mice (similar to cell culture studies). This group is called LG + Ins. The mice were fed normal chow diet after two hours of post insulin injection. This process was repeated for two weeks.

The control (C) mice groups were on continuous feed. The third group (C + Ins) was on continuous feed with insulin injection at 10 am (time same as LG + Ins). The fourth group was on fasting (similar to LG + Ins group), without any insulin injection. This group was kept to understand the effect of fasting alone on insulin signaling of peripheral tissues in mice. Mice in all groups had free access to water all the time (Fig. S3A). We have given insulin injection of final concentration of 8.6nM in all mice. We have used similar body weight mice.

### Insulin ELISA Assay

The plasma was isolated from whole blood in the presence of EDTA and circulating levels of insulin in blood were determined using insulin ELISA kit (Merck #EZRMI-13K).

### Glucose tolerance test

Following overnight fasting, mice were given an intraperitoneal dose of glucose (3g/kg body weight). Blood glucose levels were assayed at 0, 30, 60, 90 and 120 min post injection. The blood samples were collected from tail-veins of conscious mice. Blood glucose was measured using the ACCU-CHEK® glucometer (Roche).

### Insulin tolerance test

Following 2 hours of fasting, mice were given an intraperitoneal dose of insulin (0.25 U/kg body weight). Blood glucose levels were assayed at 0, 10, 20 and 30 min post injection.

### Immunohistopathological of mouse tissues

Formalin fixed samples of liver, skeletal muscle white adipose tissues and pancreatic islets were outsourced. Tissue processing was done to dehydrate in ascending grades of alcohol, clearing in xylene and embedded in paraffin wax. Paraffin wax embedded tissue blocks were sections at 3-4 μm thickness with the Rotary Microtome. All the slides of pancreas were stained with Hematoxylin & Eosin (H & E) stain. The prepared slides were examined under microscope by Pathologist to note histopathological lesions, if any.

### Statistics

The statistical significance of the data was assessed by t-test and one-way anova using mean ± standard deviation in graphpad prism 8 version. Statistical data significance levels were represented as *P*<0.05 (*), *P*<0.01 (**) and *P*<0.001 (***).

## Acknowledgments

The authors thank NCCS core facility and animal house facility. We thank Dr. Jomon Joseph for his valuable suggestions. We thank Bionivid and Nucleome to provide bioinformatic support.

## Author Contributions

SB contributed to study design, information interpretation, performed all the experiments and wrote the first draft of the manuscript. SM and AK contributed to portion of the experiment. DM and DD contributed to bioinformatic analysis. MKB provided stable cell line CHO-GLUT4 cells. VS conceptualized the experiments, supervised the project and manuscript editing.

## Data Availability

The datasets generated in this study are deposited into European Nucleotide Archive. The BRL-3A samples accession no. ERR7929863 to ERR7929866). The CHO-GLUT4 samples were submitted to ENA (accession no. are ERR8006957 and ERR8006957). This article contains supplementary data.

